# Butyrate, valerate, and niacin ameliorate anaphylaxis by suppressing IgE-dependent mast cell activation: Roles of GPR109A, PGE_2_, and epigenetic regulation

**DOI:** 10.1101/2023.02.19.529168

**Authors:** Kazuki Nagata, Daisuke Ando, Tsubasa Ashikari, Kandai Ito, Ryosuke Miura, Izumi Fujigaki, Miki Ando, Naoto Ito, Hibiki Kawazoe, Yuki Iizuka, Mariko Inoue, Takuya Yashiro, Masakazu Hachisu, Kazumi Kasakura, Chiharu Nishiyama

## Abstract

Short chain fatty acids (SCFAs) were recently shown to modulate the development and functions of immune-related cells. However, the molecular mechanisms by which SCFAs regulate mast cells (MCs) are not fully understood. We found that the oral administration of valerate or butyrate ameliorated passive systemic anaphylaxis in mice. Butyrate and valerate suppressed the IgE-mediated degranulation of bone marrow-derived MCs, which were eliminated by pertussis toxin and by the knockdown of *Gpr109a*. A treatment with trichostatin A suppressed IgE-mediated MC activation and reduced the surface expression level of FcεRI on MCs. Acetylsalicylic acid and indomethacin attenuated the suppressive effects of SCFAs on degranulation. The degranulation degree was significantly decreased by the treatment with PGE_2_ whose release from MCs was markedly enhanced by SCFAs. The SCFA-mediated amelioration of anaphylaxis was exacerbated by COX inhibitors and an EP3 antagonist. The administration of niacin, a ligand of GPR109A, alleviated the symptoms of passive cutaneous anaphylaxis, which was inhibited by COX inhibitors and the EP3 antagonist.

**Key Messages:** Short chain fatty acids (SCFAs), particularly butyrate and valerate, suppress the IgE-mediated activation of mast cells (MCs) *in vivo* and *in vitro*.

SCFAs enhance the release of PGE_2_ from MCs, which inhibits the IgE-mediated activation of MCs.

Niacin, a ligand of GPR109A, ameliorates IgE-dependent anaphylaxis.

The administration of COX inhibitors or an antagonist of PGE_2_ receptor 3 (EP3) inhibited the suppressive effects of butyrate and niacin on IgE-dependent anaphylaxis.

## Introduction

Dietary fibers that are barely digested in the mammalian digestive system are fermented by the bacteria that constitute the intestinal flora. Short chain fatty acids (SCFAs), including acetate, butyrate, propionate, and valerate, are produced in the colon during the fermentation process as secondary metabolites. The effects of intestinal SCFAs on immune responses have recently become topics of interest. Previous studies indicated that SCFAs contributed to the maintenance of homeostasis by modulating the development and function of immune-related cells ^1, 2^. SCFAs have been shown to exert beneficial effects on immune-related diseases. Mice kept under germ-free conditions or fed a low-fiber diet showed an exacerbated pathology and symptoms of inflammatory bowel disease, airway hypersensitivity, experimental autoimmune encephalomyelitis, and rheumatoid arthritis due to a deficiency in SCFAs ^3–8^. SCFAs attenuate these inflammatory diseases by accelerating Treg development ^4, 5, 7^, increasing IL-10 production by lymphoid cells ^4, 8^, suppressing the activation of neutrophils and eosinophils ^3^, and regulating T cell development via the modulation of monocyte functions ^6^. Intestinal epithelial cells and immune cells take up SCFAs through monocarboxylate transporters and/or sense SCFA signals via G protein-coupled receptors (GPCRs) on the cell surface. SCFAs exhibit an additional function as HDAC inhibitors, inducing the expression of genes involved in the anti-inflammatory function of immune cells.

Mast cells (MCs), which develop in bone marrow and terminally differentiate in the mucosa and connective tissues of the whole body, play a key role in IgE-mediated allergic diseases, such as pollinosis and food allergy. The cross-linking of the high-affinity receptor for IgE, FcεRI, by the IgE and antigen (Ag) complex induces rapid degranulation, eicosanoid release, and cytokine production in MCs, resulting in allergic symptoms. Although accumulating evidence recently showed the anti-allergic effects of SCFAs targeting MCs ^9–11^, the molecular mechanisms by which SCFAs modulate MC functions remain unknown. In the present study, we demonstrated that the oral administration of butyrate and valerate ameliorated IgE-mediated anaphylaxis in mice, and that SCFAs significantly reduced the degree of IgE-induced degranulation and cytokine production of bone marrow-derived MCs (BMMCs). We also investigated the molecular mechanisms underlying the suppressive effects of SCFAs *in vitro* and *in vivo*, by examining the roles of GPR109A, HDAC inhibition (HDACi) activity, the NRF2 pathway, and prostaglandins (PGs) in the effects of SCFAs on MCs. Based on the present results, we concluded that butyrate and valerate regulated MCs via GPR109A and by the HDACi activity, with the accelerated synthesis of PGE_2_, resulting in the amelioration of anaphylaxis.

## Results

### Orally administered butyrate and valerate suppress IgE-mediated PSA

To evaluate the efficacy of SCFAs to prevent MC-mediated allergic responses *in vivo*, we utilized the PSA model, and found that the decrease in body temperature caused by the IgE-mediated activation of MCs was significantly abated by the administration of butyrate (Fig. 1A) and valerate (Fig. 1B). To confirm whether the intake of SCFAs under this experimental condition affected the development of Treg, which is known to be enhanced by SCFAs, we examined the frequency of Treg cells in the mesenteric lymph node (MLN) and spleen. In the mice that received 100 μmol butyrate or valerate once per day for 4 to 6 days, an increase in Treg cells was not observed in the MLN or spleen (Figs. 1C, 1D). Footpad swelling following a subcutaneous (s.c.) injection of IgE and i.v. injection of Ag was also reduced in mice that received orally administered butyrate (Fig. 1E), suggesting that the uptake of SCFAs via the digestive tract suppressed not only systemic anaphylaxis, but also peripheral cutaneous anaphylaxis. These results indicate that butyrate and valerate exerted suppressive effects on the MC-mediated allergic responses *in vivo* independent of Treg development.

**Fig. 1.**
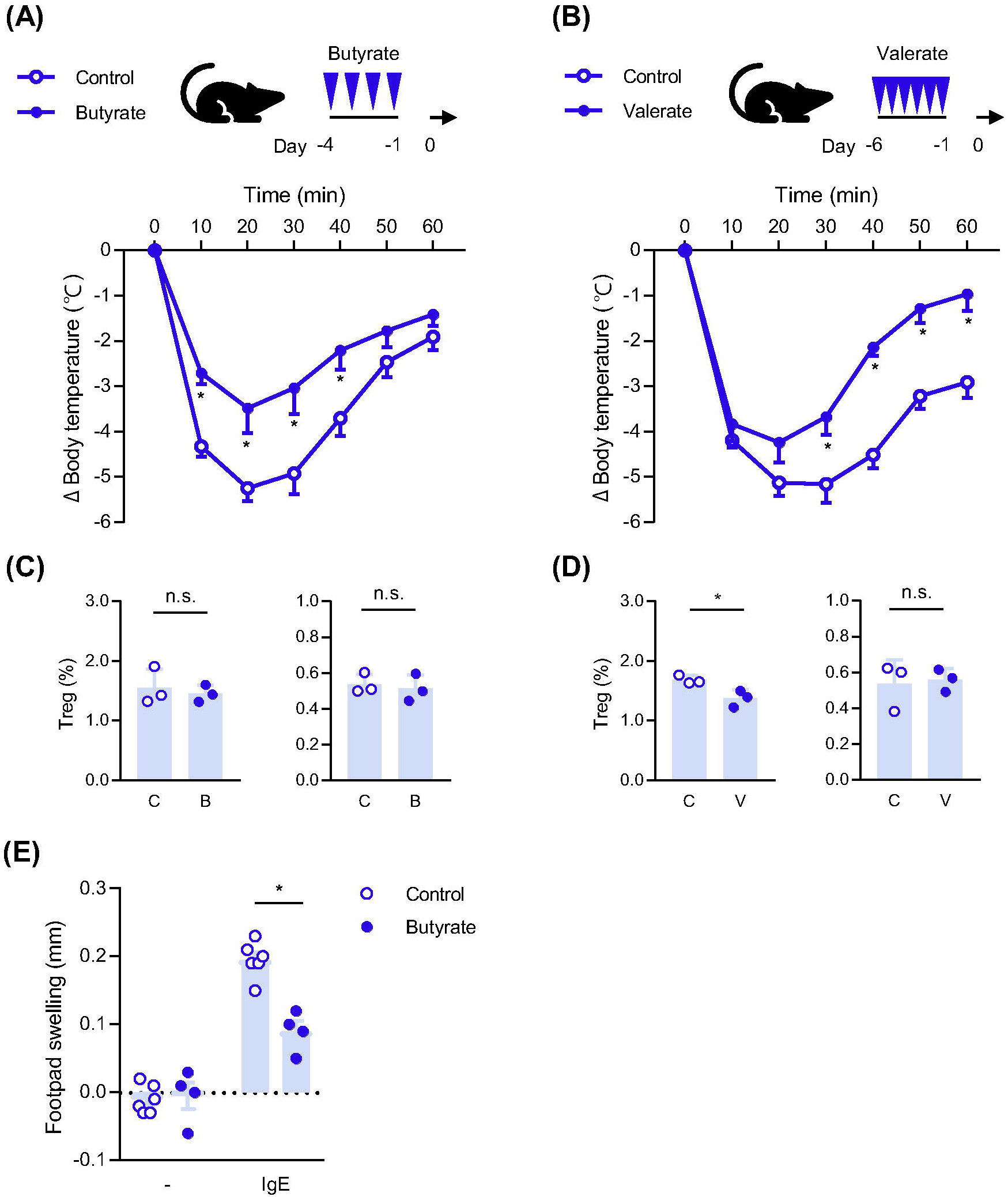
Butyrate and valerate alleviate the IgE-induced anaphylaxis reaction. (**A**) Changes in the body temperature of control and butyrate-administered mice following PSA. Mice were orally administered 100 μmol (200 μl of 500 mM) butyrate or saline once a day for 4 days prior to PSA. Open circle, control (n = 10); closed circle, butyrate administration (n = 9). Data were pooled from 2 independent experiments. (**B**) Changes in the body temperature of control and valerate-administered mice following PSA. Mice were orally administered 100 μ mol (200 μl of 500 mM) valerate or ethanol-PEG solution once a day for 6 days prior to PSA. Open circle, control (n = 6); closed circle, valerate administration (n = 5). Data were pooled from 2 independent experiments. (**C**, **D**) Frequency of Foxp3^+^ cells in CD4^+^ T cells isolated from the MLN (left) and spleen (right) of mice administered butyrate (B) or saline as the control of butyrate (C) on the same schedule as in Figure 2A (**C**), or mice administered valerate (V) or ethanol-PEG as the control (C) on the same schedule as in Figure 2B (**D**). (**E**) Anaphylaxis induced on the footpad by the s.c. injection of IgE (right footpad) or vehicle (left footpad) and a subsequent i.v. injection of Ag. Footpad thickness was measured before and after the Ag injection. Footpad swelling (mm) was calculated as follows. (Footpad swelling) = (Footpad thickness after the Ag injection) – (Footpad thickness before the Ag injection) Sidak’s multiple comparison tests (**A**, **B**, **E**), and a two-tailed Student’s *t*-test (**C**, **D**) were used for statistical analyses. *, *p*<0.05

### Effects of butyrate and valerate on the degranulation and cytokine production in IgE-stimulated MCs

BMMCs pretreated with various SCFAs were stimulated with IgE and Ag to evaluate the effects of SCFAs on MC activation. The activity of β-hexosaminidase released from MCs was assessed as an index of the immediate response level (Fig. 2A). As shown in Fig. 2B, the frequency of DAPI-stained cells remained unchanged in the presence of 10 mM SCFAs, suggesting that SCFAs did not induce apparent toxicity. Under this experimental condition, five out of the six SCFAs examined, namely, propionate, butyrate, valerate, isobutyrate, and isovalerate, suppressed the degranulation of IgE-stimulated BMMCs in a dose-dependent manner (Fig. 2A). The suppressive effects of these SCFAs at 1 mM were significant, whereas acetate did not affect the degree of degranulation even at 10 mM (Fig.2A). These results suggest that these five SCFAs suppressed the IgE-induced degranulation of MCs. We also examined sodium butyrate to exclude the effects of pH changes caused by the acids, and confirmed that sodium butyrate significantly reduced IgE-stimulated degranulation as well as butyrate (Fig. 2C).

**Fig. 2.**
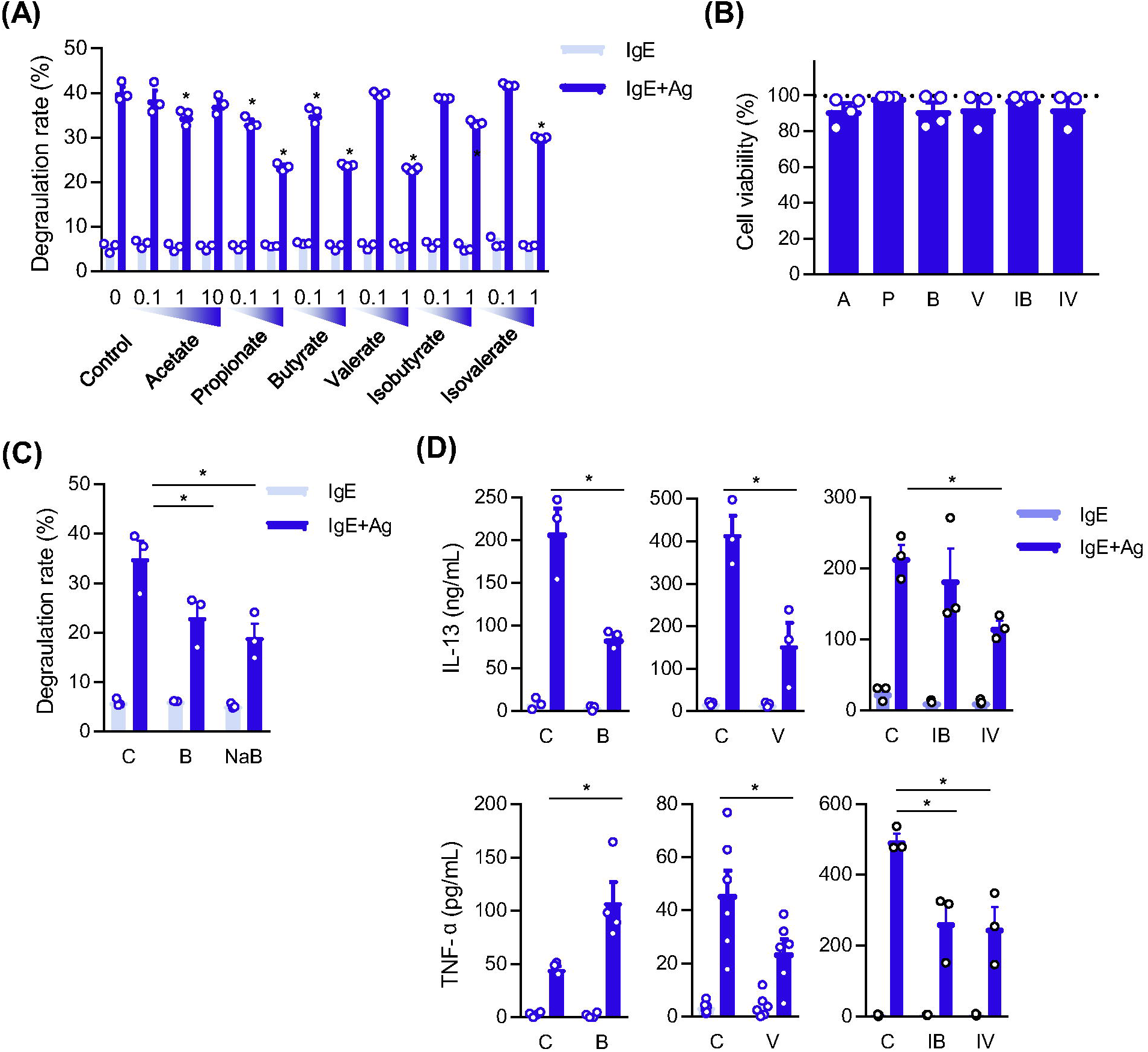
SCFAs suppress the degranulation of IgE-stimulated MCs. (**A**) Effects of SCFAs on the degranulation degree of IgE-stimulated BMMCs. BMMCs were treated with the indicated concentrations of SCFAs for 48 h. Data represent the mean ± SD of triplicate samples. A typical result is shown, and representative results were obtained from 3 independent experiments. (**B**) Viability of BMMCs incubated in the presence of 10 mM SCFAs for 48 h. A; acetate, P; propionate, B; butyrate, V; valerate, IB; isobutyrate, IV; isovalerate. Survival rates were assessed by DAPI staining using flow cytometry. Data represent the mean ± SD of 3 independent experiments. (**C**) Degranulation degree of IgE-stimulated BMMCs pretreated with 1 mM butyrate (B), 1mM sodium butyrate (NaB), or without SCFA (C; control). The data represent the mean ± SEM of 3 independent experiments. (**D**) Effects of SCFAs on the amount of cytokines produced by IgE-stimulated BMMCs. Data represent the mean ± SEM of 3 independent experiments. C; control, B; butyrate, V; valerate, IB; isobutyrate, IV; isovalerate. A two-tailed Student’s t-test (left and middle in **D**) and Dunnett’s multiple comparison tests (**A**, **B**, **C**, right in **D**) were used for statistical analyses. *, *p*<0.05.

The effects of SCFAs on cytokine production by MCs was examined. As shown in Fig. 2D, IL-13 release from IgE-stimulated MCs was significantly suppressed by the pretreatment with butyrate, valerate, and isovalerate, and was slightly suppressed by the isobutyrate treatment. The IgE-induced production of TNF-α was significantly inhibited by the treatment with valerate, isobutyrate, and isovalerate, whereas the butyrate treatment promoted the production of TNF-α.

The IgE-induced activation of MCs was initiated by the binding of IgE to FcεRI. The expression level of FcεRI is associated with the degree of degranulation and is a risk factor for allergic diseases ^12–15^. To evaluate the effects of SCFAs on FcεRI expression, we performed a flow cytometric analysis and found that FcεRI levels on the cell surface were decreased by the pretreatment with butyrate and valerate (Fig. 3A). FcεRI comprises 3 subunits, namely, α, β, and γ, and the transcription factors PU.1, GATA1, and GATA2 regulate the cell type-specific expression of α and β ^16–20^. To clarify whether the down-regulation of surface FcεRI occured in a transcription-dependent manner, we measured the mRNA levels of FcεRI subunits (Fig. 3B) and related transcription factors (Fig. 3C) by quantitative PCR. The obtained results revealed that butyrate and valerate did not reduce but rather tend to increase the mRNA levels of *Fcer1a*, *Ms4a2*, *FceR1g*, *Spi1*, *Gata1*, and *Gata2*. Therefore, valerate and butyrate ma have reduced FcεRI levels on the cell surface without inhibiting the transcription of FcεRI subunit genes.

**Fig. 3.**
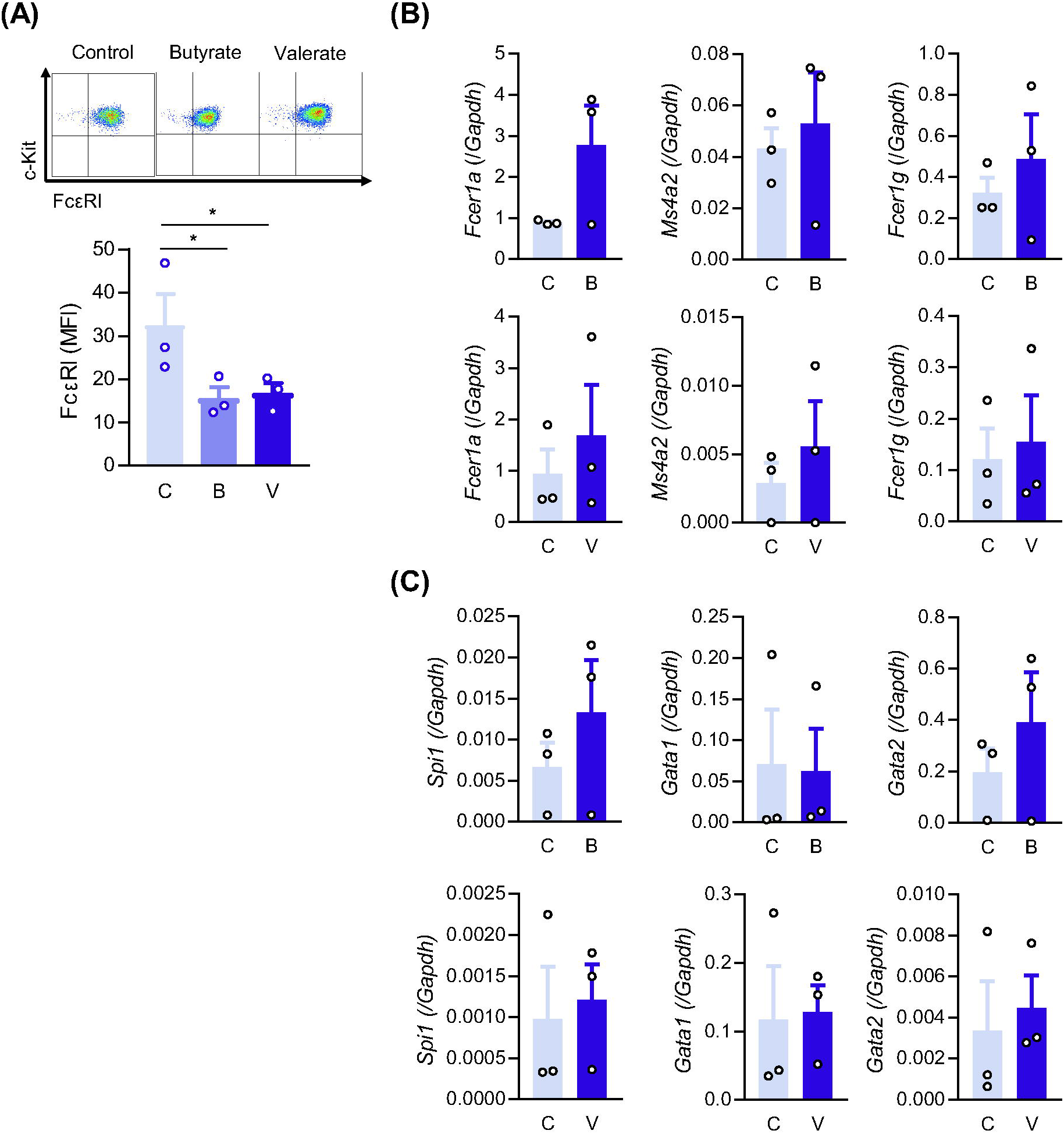
Effects of butyrate and valerate on the cell surface expression level of FcεRI and on the mRNA levels of FcεRI subunits. (**A**) Cell surface expression levels of FcεRI and c-kit on SCFA-treated BMMCs. A typical flow cytometry profile (top) and relative FcεRI expression level based on data obtained from 3 independent experiments (bottom). C; control, B; butyrate, V; valerate. Dunnett’s multiple comparison test was used for statistical analysis. *, *p*<0.05. (**B**, **C**) mRNA levels of FcεRI subunits (**B**) and FcεRI-related transcription factors (**C**) in BMMCs treated with butyrate, or valerate. *Fcer1a* (encoding FcεRI α subunit), *Ms4a2* (FcεRI β), *Fcer1g* (FcεRI γ), *Spi1* (PU.1), *Gata1* (GATA1), and *Gata2* (GATA2). BMMCs incubated in the presence of 1 mM butyrate (B), or valerate (V) for 48 h were harvested to measure mRNA levels. Tukey’s multiple comparison test was used. *, *p*<0.05 versus the control without the SCFA treatment (C).

These results indicate that butyrate and valerate suppressed the IgE-mediated degranulation and cytokine release of MCs, and also that the decrease observed in the cell surface level of FcεRI was partly involved in the suppressive effects of SCFAs on the IgE-mediated activation of MCs.

### Butyrate and valerate suppress the activation of MCs via GPR109A

To elucidate the molecular mechanisms by which butyrate and valerate modify the function of MCs, we examined cell surface molecules in order to identify a candidate transporter and/or receptor for SCFA. A quantitative PCR analysis showed that BMMCs expressed detectable amounts of mRNAs for the solute carrier group of membrane transport proteins (*Slc5a8*, *Slc5a12*, and *Slc16a1*) and GPCRs (*Gpr41*, *Gpr43*, and *Gpr109a*). Moreover, the mRNA expression levels of *Slc16a1* and *Gpr109a* were higher than those of other mRNAs (Figs. 4A, 4B). To confirm the involvement of the transporter and receptor in SCFA signaling, we pretreated BMMCs with a reagent that inhibits the transporter or receptor for 1 h prior to the addition of SCFAs. As shown in Fig. 4C, the presence of 2-cyano-4-hydroxyphenyl acrylic acid (monocarboxylate transporter inhibitor) did not affect the SCFA-mediated suppression of degranulation. In contrast, PTX, an inhibitor of Gi/o proteins, counteracted the suppressive effects of butyrate and valerate (Fig. 4D), suggesting that Gi/o-type GPCR functions are required for the butyrate- and valerate-mediated suppression of MC activation. We also found that mRNAs for GPR43 and GPR109A were expressed in human MCs (Fig. 4E). To clarify the involvement of GPR109A, which was a Gi-GPCR that was expressed at higher levels than other Gi-GPCRs (GPR41 and GPR43) in mouse and human MCs (Figs. 4B, 4E), we performed a knockdown experiment using siRNA. When butyrate and valerate significantly suppressed the degranulation of control siRNA-introduced BMMCs, *Gpr109a* siRNA transfectants in which *Gpr109a* mRNA was effectively knocked down (Fig. S1A) exhibited markedly enhanced degranulation (Fig. 4F). This result supports the hypothesis that GPR109A is a receptor for butyrate and valerate on MCs. The increase in degranulation by the knockdown of *Gpr109* was more restrictive for butyrate-treated MCs than for valerate-treated MCs, suggesting that the effects of butyrate were partly dependent on GPR109A. The knockdown of *Gpr109a* up-regulated the degranulation of non-treated control BMMCs. This result suggests that GPR109A signaling constitutively suppressed MC activation. Therefore, to clarify the role of GPR109A in the IgE-dependent activation of MCs, we conducted a GPR109A overexpression experiment. When GPR109A was constitutively overexpressed in BMMCs using a retroviral vector (Fig. S1B), the suppressive effects of SCFAs were enhanced and the degree of degranulation was markedly reduced even in the absence of SCFAs (Fig. 4G).

**Fig. 4.**
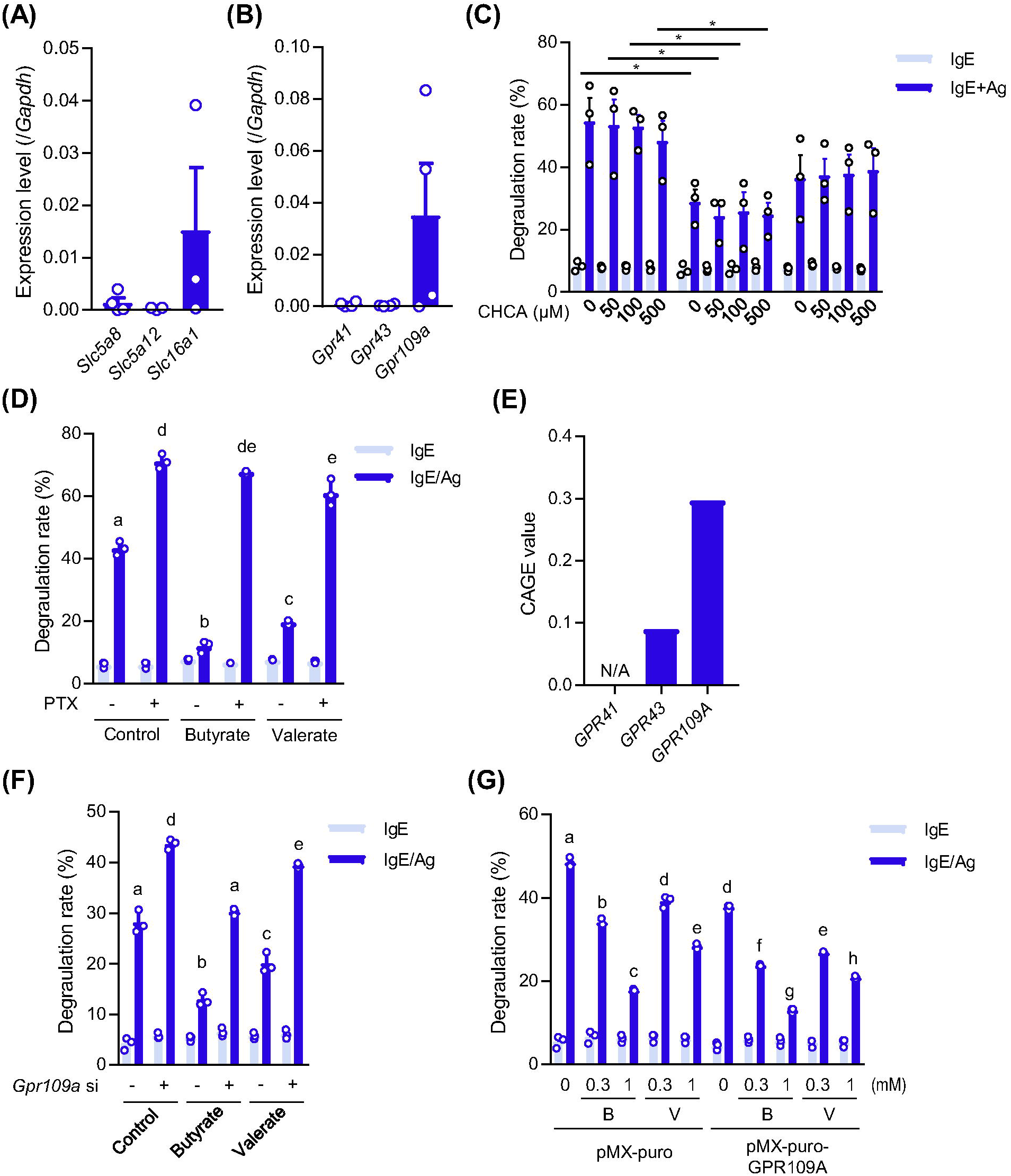
Involvement of GPR109A in the butyrate- and valerate-mediated suppression of degranulation. (**A, B**) Expression levels of mRNAs for transporters (**A**) and GPCRs (**B**) in MCs. Data represent the mean ± SEM of 3 (*Slcs*) and 4 (*Gprs*) independent experiments. (**C**) Degranulation degree of BMMCs treated with the transporter inhibitor, CHCA, in the presence and absence of 1 mM SCFAs. The indicated concentrations of CHCA were added to the culture media 1 h before the SCFA treatment. Data represent the mean ± SEM of 3 independent experiments. (**D**)Effects of 0.1 μg/mL of PTX, a Gi protein inhibitor on SCFA-dependent suppression. Data represent the mean ± SD of triplicate samples, and representative results were obtained from 3 independent experiments. (**E**)Expression levels of mRNAs for GPCRs in human MCs. Data were obtained from “processed expression data of all samples for CAGE human PRJDB1099 (FANTOM5)” (https://figshare.com/articles/dataset/RefEx_expression_CAGE_all_human_FANTOM5_tsv_zip/4028613). (**F**) Degree of degranulation in MCs with the knockdown of GPR109A. Data represent the mean ± SD of triplicate samples, and the representative results were obtained from 3 independent experiments. (**G**) Degranulation degree of GPR109A-overexpressing and control (mock) BMMCs. GPR109A-overexpressing and control BMMCs were stimulated with IgE plus Ag after a pre-incubated in the presence or absence of the indicated concentrations of SCFAs. Data represent the mean ± SD of triplicate samples, and representative results were obtained from 3 independent experiments. Dunnett’s multiple comparison test (**C**) and Tukey’s multiple comparison test (**D**, **F**, and **G**) were used. *, *p*<0.05.

Based on these results, we concluded that GPR109A was involved in the suppression of the sIgE-dependent degranulation of MCs and that valerate and butyrate functioned as ligands for GPR109A.

### Trichostatin A (TSA)-treatment suppresses the IgE-mediated activation of MCs

SCFAs exhibit HDACi activity, which enhances the anti-inflammatory functions of immune cells by inducing the expression of genes including Foxp3 and IL-10 ^5, 8^. The order of anti-allergic effects in Fig. 2A (butyrate > valerate >> acetate) is consistent with the order of HDACi activity among SCFAs ^8^. To evaluate the effects of HDACi activity on the IgE-mediated activation of MCs, we analyzed MCs that were treated with TSA, an inhibitor of class I and II HDACs. The pretreatment of MCs with TSA (5 – 20 nM) for 24 h before the IgE-mediated stimulation significantly and dose-dependently suppressed degranulation (Fig. 5A) without affecting the frequency of DAPI-stained dead cells (Fig. 5B). The release of IL-13 and TNF-α from IgE-stimulated MCs was significantly inhibited by the pretreatment with 20 nM TSA (Fig. 5C).

**Fig. 5.**
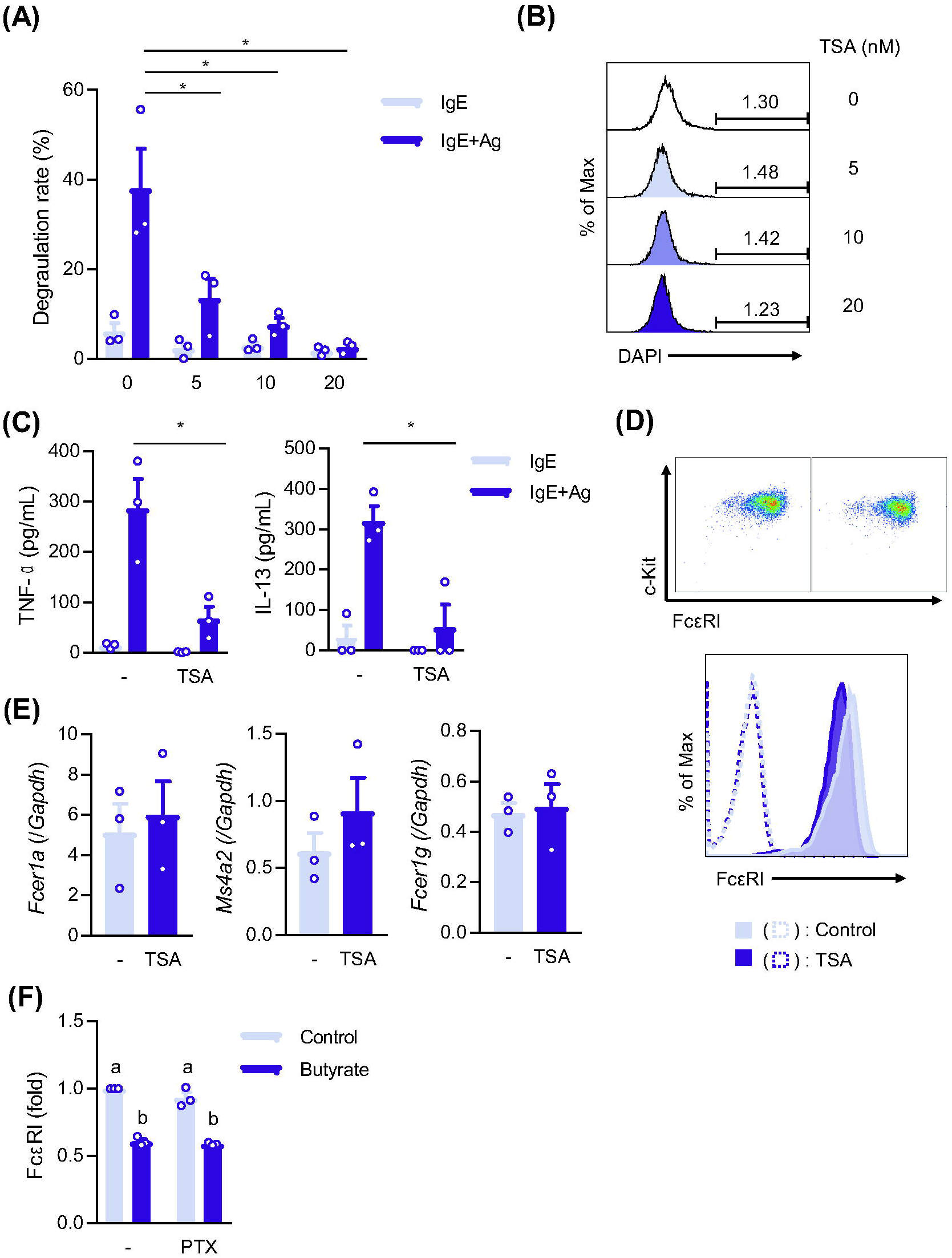
Effects of TSA on the IgE-mediated activation of and FcεRI levels in MCs. (**A**) IgE-mediated degranulation of TSA-treated BMMCs. BMMCs were preincubated with the indicated concentrations (nM) of TSA for 24 h. Data represent the mean ± SEM of 3 independent experiments. (**B**) Percentage of DAPI-stained cells in BMMCs pretreated with the indicated concentrations of TSA for 24 h. (**C**) IgE-induced cytokine release from TSA-treated BMMCs. BMMCs incubated in the presence or absence of 20 nM TSA for 24 h were stimulated with IgE plus Ag, and culture supernatants at 3 h after the stimulation were harvested to measure the concentrations of TNF-α and IL-13. Data represent the mean ± SEM of 3 independent experiments. (**D**, **E**) Surface expression levels of FcεRI (**D**) and mRNA levels of *Fcer1a*, *Ms4a2*, and *Fcer1g* (**E**) in BMMCs treated with 20 nM TSA for 24 h. Data represent the mean ± SEM of 3 independent experiments. (**F**) FcεRI expression levels on BMMCs, which were pretreated in the presence or absence of 0.1 μg/mL PTX for 1 h and further incubated with or without 1 mM butyrate for 48 h. Dunnett’s multiple test (A), a *t*-test (C, E), and Tukey’s multiple comparison test (F) were used. *, *p*<0.05.

The TSA treatment reduced the surface expression levels of FcεRI (Fig. 5D), even though the mRNA levels of FcεRIα, β, and γ were not decreased in TSA-treated MCs (Fig. 5E), which is consistent with the results obtained from butyrate-treated MCs (Fig. 2). We also found that the suppressive effects of butyrate on FcεRI expression levels in MCs were not inhibited by the PTX treatment (Fig. 5F).

These results suggest that SCFA-induced reductions in the cell surface expression level of FcεRI were mediated by the HDAC inhibition rather than by the Gi-GPCR-stimulation.

### NRF2 is activated by SCFA, but is not involved in suppressive effect of SCFAs

Butyrate has been shown to activate the transcriptional regulator NRF2, which alleviates oxidative stress, in various cells ^21–23^. Furthermore, the activation of NRF2 is frequently observed in MCs treated with anti-allergic compounds ^24–26^. Hmox1, a target gene of NRF2, exhibited the greatest increase in its mRNA level following the treatment of human MCs with butyrate ^9^, however, the role of NRF2 in human MCs was not analyzed. Based on these findings, we investigated whether NRF2 was activated in SCFA-treated MCs and played a role in the suppressive effects of SCFAs on MCs. As shown in Figs. 6A and 6B, the mRNA levels of Hmox1 and Nrf2 were elevated in butyrate-treated BMMCs, and these increases were sustained for at least 48 h under our experimental conditions. In addition, we found that Hmox1 mRNA levels, which were increased by the IgE-mediated stimulation, were further up-regulated in the presence of butyrate (Fig. 6C), and that the TSA treatment increased Hmox1 mRNA levels in MCs (Fig. 6D). To clarify whether the SCFA-induced activation of NRF2 in MCs was involved in the suppression of IgE-mediated activation, we conducted an experiment using an NRF2 inhibitor and found that the inhibition of the NRF2 pathway did not reduce the suppressive effects of butyrate on the degranulation and cytokine release (Fig. 6E).

**Fig. 6.**
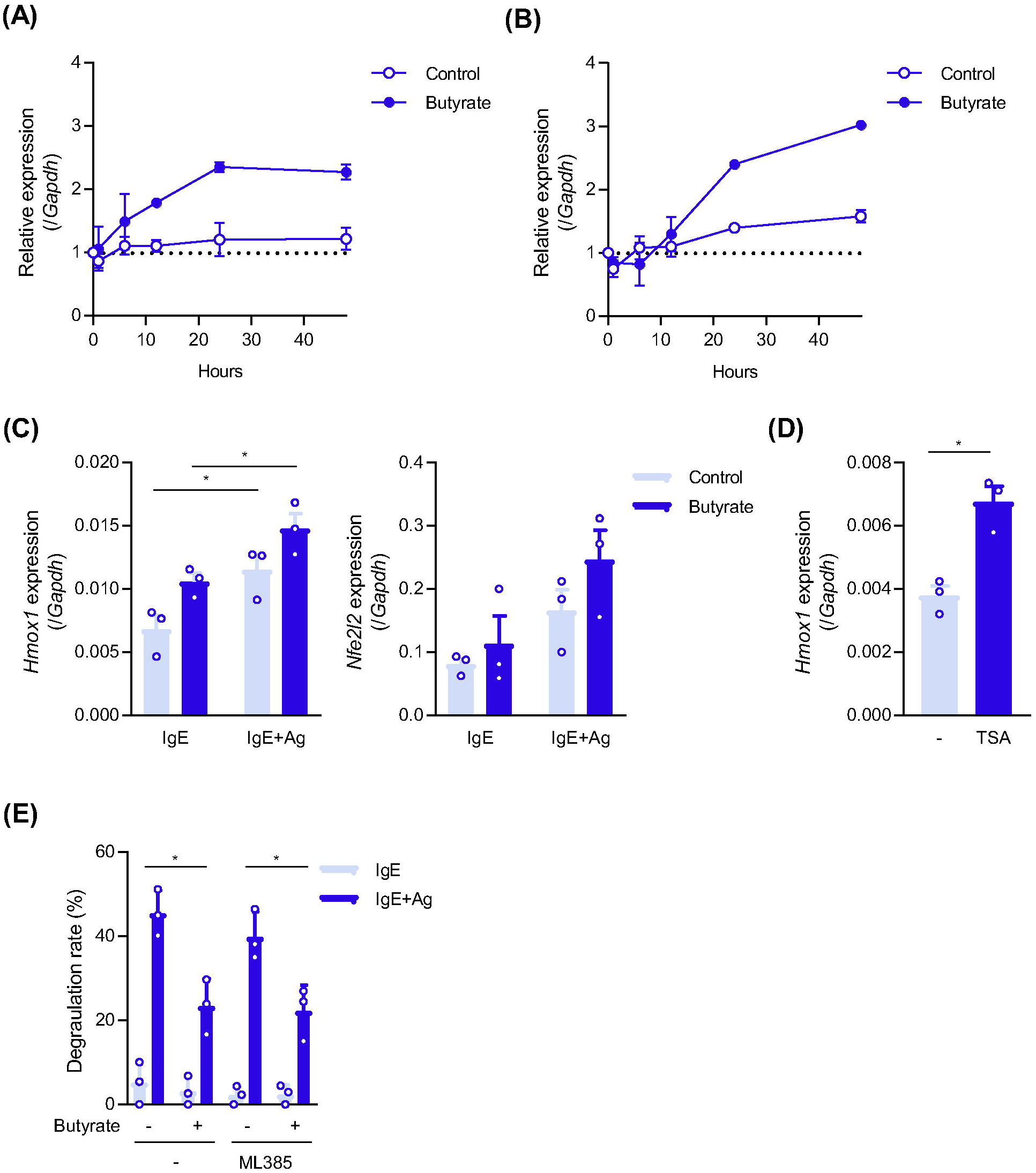
SCFA-induced activation of the NRF2 pathway in MCs. (**A**, **B**) Effects of butyrate on the transcription levels of NRF2 pathway-related genes. The time-course of the relative mRNA levels of *Hmox1* (encoding HO-1) (**A**) and *Nfe2l2* (NRF2) (**B**) in butyrate-treated BMMCs. (**C**) mRNA levels of *Hmox1* (left) and *Nfe2l2* (right) in butyrate-treated and/or IgE-stimulated BMMCs. BMMCs pretreated with 1 mM butyrate for 48 h were stimulated with IgE plus Ag, and then harvested after an additional incubation for 1 h. (**D**) mRNA levels of *Hmox1* in BMMCs treated with or without 20 nM TSA for 24 h. (**E**) The butyrate-mediated suppression of degranulation was not affected by an NRF2 inhibitor. BMMCs were pretreated with 1 mM butyrate and 5 μM ML385 (an NRF2 inhibitor) for 48 h before the IgE-induced degranulation assay. Data represent the mean ± SEM of 2 (**A**, **B**) and 3 (**C**, **D**, **E**) independent experiments. Tukey’s multiple comparison test (**C**, and **E**), and a *t*-test (**D**) were used for statistical analyses. *, *p*<0.05.

These results indicate that the activation of the NRF2-HO-1 pathway was not involved in the suppression of IgE-mediated activation in SCFA-treated MCs; however, SCFAs and TSA activated this pathway in MCs.

### Butyrate and valerate increase the release of PGE_2_, which suppresses degranulation

GPR109A is classically known as a receptor for niacin/nicotinic acid/vitamin B3. Moreover, the intake of excess amounts of niacin induces a niacin flash due to the enhanced release of PGs. Based on these findings, we hypothesized that the treatment of MCs with butyrate and valerate may induce the production of PGs, which affects the activation degree of MCs. To evaluate the roles of PGs in MC activation, we analyzed the degranulation degree of SCFA-treated MCs in the presence of the non-steroidal anti-inflammatory drugs (NSAIDs), ASA and indomethacin. The suppression of degranulation in valerate-treated MCs was counteracted by the addition of ASA, whereas the effects of ASA and indomethacin on butyrate-induced suppression were moderate (Figs. 7A, 7B).

**Fig. 7.**
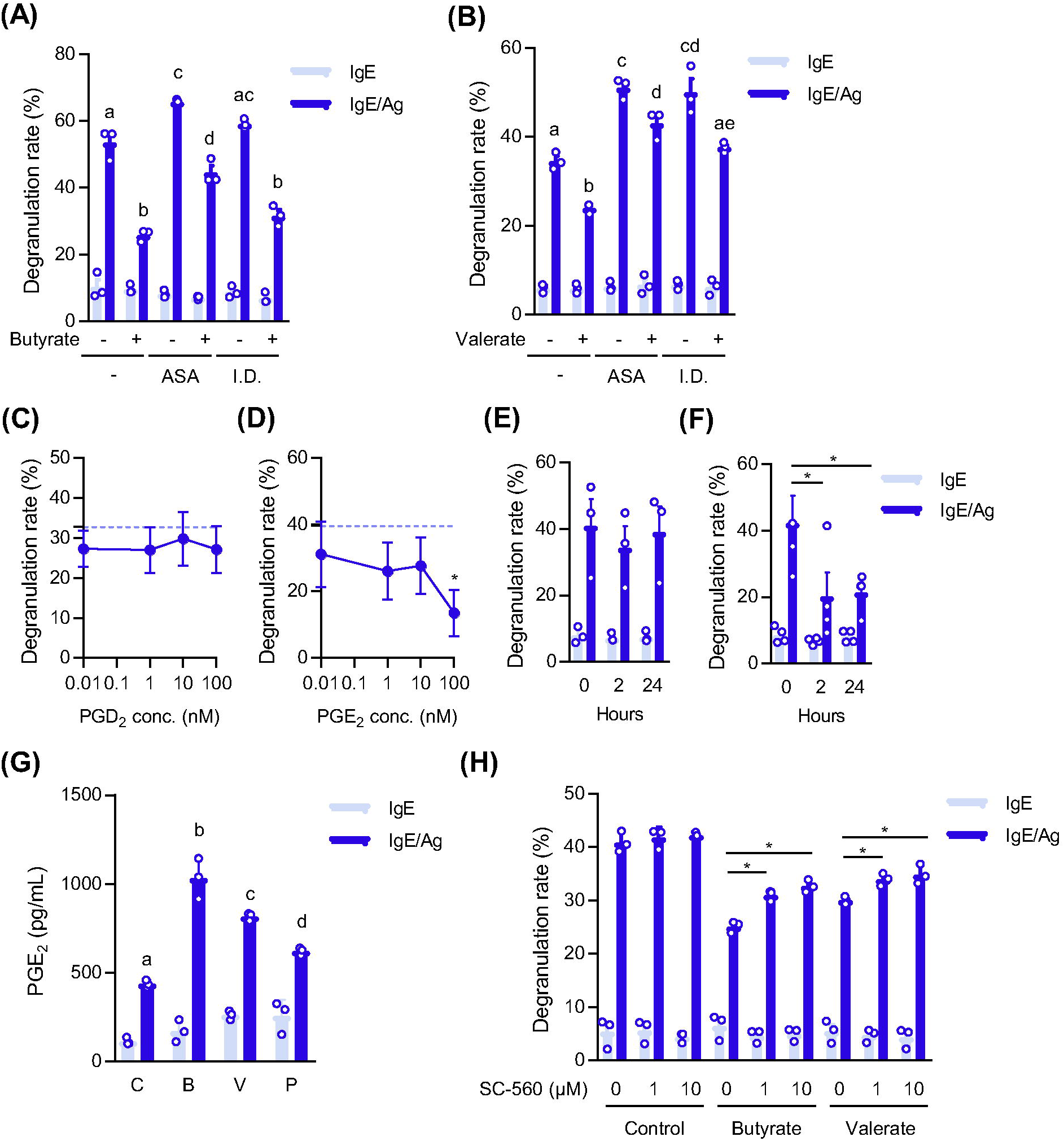
Roles of COX and PGs in suppressive effects of butyrate and valerate. (**A**, **B**) Effects of the ASA and indomethacin pretreatment on the butyrate (**A**)- or valerate (**B**)-mediated suppression of degranulation. Data represent the mean ± SD of triplicate samples and the representative results were obtained from 3 independent experiments. (**C**, **D**) Effects of PGD_2_ and PGE_2_ on the degranulation of MCs. BMMCs were pretreated with the indicated concentrations of PGD_2_ (**C**) or PGE_2_ (**D**) for 24 h prior to the degranulation assay. Data represent the mean ± SEM of 3 independent experiments. (**E**, **F**) Degranulation degree of MCs pretreated with PGs. A total of 100 nM PGD_2_ (**E**) or PGE_2_ (**F**) was added to the culture media 2 h or 24 h before IgE stimulation. Data represent the mean ± SEM of 3 (PGD_2_) and 4 (PGE_2_) independent experiments. (**G**) SCFAs up-regulate the amount of PGE_2_ released from MCs. Data represent the mean ± SD of triplicate samples, and representative results were obtained in another independent experiment. (**H**) Effects of the inhibitor of COX-1, SC-560, on the SCFA-mediated suppression of MC degranulation. BMMCs pretreated with the indicated concentrations of SC-560 for 2 h were incubated for an additional 48 h in the presence or absence of 1 mM of SCFAs. Data represent the mean ± SD of triplicate samples, and representative results were obtained from 3 independent experiments. Tukey’s multiple comparison test (**A**, **B**, **G**, **H**) and Dunnett’s multiple comparison test (**C**, **D**, **E**, **F**) were used for statistical analyses. *, *p*<0.05.

PGD_2_ and PGE_2_ are preferentially produced by activated MCs. Therefore, we evaluated the effects of PGD_2_ and PGE_2_ on the degranulation levels. Although pretreatment with 1-100 nM PGD_2_ for 24 h did not affect the degranulation of MCs (Fig. 7C), that with PGE_2_ exerted inhibitory effects (Fig. 7D). Similar suppressive effects of PGE_2_ were observed following its addition 2 h prior to the stimulation (Fig. 7F), whereas PGD_2_ did not inhibit degranulation at either time point (Fig. 7E). The measurement of PGE_2_ concentrations in culture media revealed that the IgE stimulation strongly promoted the release of PGE_2_, which was already enhanced by butyrate, valerate, and propionate (Fig. 7G). Furthermore, SCFA-treated MCs released slightly more PGE_2_ than non-treated MCs, even in the steady state (Fig. 7G).

We recently reported that a treatment with ASA or the knockdown of cyclooxygenases (COXs), particularly COX-1, promoted the IgE-induced activation of BMMCs ^27^. To reveal the role of COX-1 in the suppression of degranulation in SCFA-treated MCs, we evaluated the effects of the COX-1-specific inhibitor SC-560 on MC degranulation. When BMMCs were pretreated with SC-560 prior to the addition of SCFAs, SC-560 attenuated the suppressive effects of SCFAs in a dose-dependent manner (Fig. 7H). Collectively, these results demonstrated that the SCFAs increased the *de novo* synthesis and release of PGE_2_, which suppressesd the activation of MCs.

### Involvement of PGs in the SCFA-mediated suppression of anaphylaxis

To investigate whether prostanoid production is involved in the effects of SCFAs *in vivo*, we performed anaphylaxis analyses of mice that were administered an NSAID. When PSA was induced in mice administered butyrate and/or indomethacin, the effects of the administration of butyrate were significantly inhibited by supplementation with indomethacin (Fig. 8A). In mice administered ethylvalerate (to avoid the corrosivity of valerate) and/or ASA, the decrease of body temperature following the Ag injection was reduced by the administration of ethylvalerate, and the suppression of PSA by ethylvalerate was counteracted by additional supplementation with ASA (Fig. 8B). The results showing that NSAIDs suppressed the SCFA-mediated amelioration of PSA suggest the involvement of PGs in the effects of SCFAs on passive anaphylaxis. To further clarify the role of PGE_2_, we conducted a PGE_2_ receptor antagonism experiment *in vivo*, and found that the administration of the EP3 antagonist inhibited the attenuating effects of butyrate on PSA (Fig. 8D) and PCA (Fig. 8E), whereas the EP4 antagonist did not affect the butyrate-mediated suppression of PCA (Fig. 8F).

**Fig. 8.**
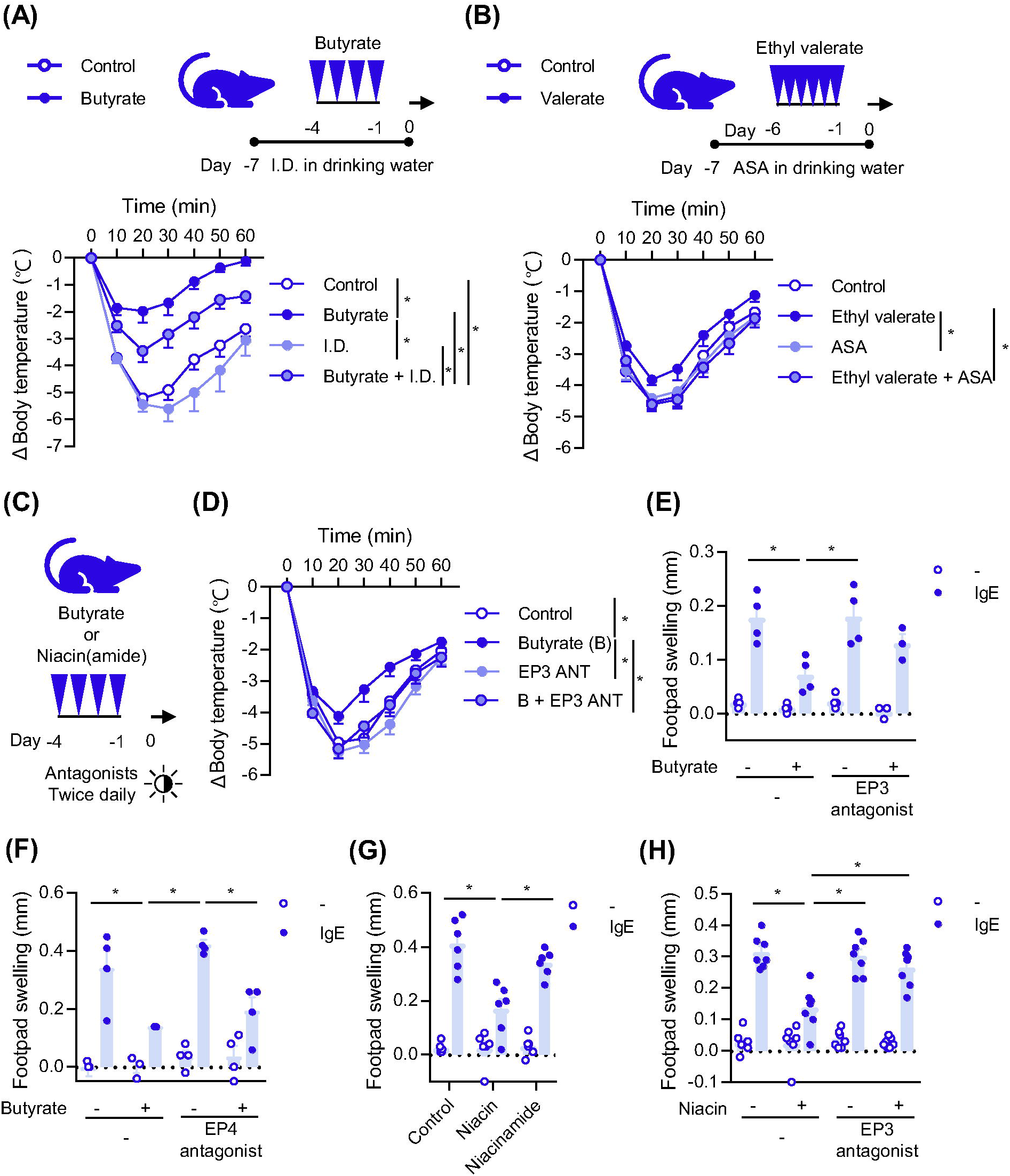
Involvement of PGs and GPR109A-signaling in the SCFA-induced suppression of anaphylaxis. (**A**) Schematics of the administration schedule of butyrate and indomethacin (I.D.) (top). Changes in body temperature after the Ag challenge (bottom). Control, the administration of saline as vehicle (n = 8); Butyrate, butyrate administration without indomethacin (n = 8); I.D., indomethacin intake without butyrate administration (n = 6); Butyrate + I.D., butyrate administration with indomethacin intake (n = 5). (**B**) Schematics of the administration schedule of ethyl valerate and ASA (top). Changes in body temperature after the Ag challenge (bottom). Control, the administration of ethanol-PEG as vehicle (n = 12); Ethyl valerate, ethyl valerate administration without ASA (n = 12); ASA, ASA intake without ethyl valerate administration (n = 12); Ethyl valerate + ASA, ethyl valerate administration with ASA intake (n = 11). (**C**) Administration schedule of EP antagonists, butyrate, and/or niacin on passive anaphylaxis in **D**-**H** of Fig. 8. (**D**) The administration of the EP3 antagonist suppressed the butyrate-mediated amelioration of PSA. Changes in body temperature after the Ag challenge for PSA. Control, vehicle (n = 12); Butyrate (B), butyrate administration without the EP3 antagonist (n = 9); EP3 ANT, the EP3 antagonist treatment without butyrate (n = 12); B+ EP3 ANT, butyrate administration and the EP3 antagonist treatment (n = 10). (**E**, **F**) Effects of the EP3 antagonist (**E**) and EP4 antagonist (**F**) on the butyrate-mediated suppression of PCA. PCA was induced in mice treated with IgE (IgE; right footpad) or vehicle (-; left footpad) by an i.v. injection of Ag. Swelling was assessed by measurements of footpad thickness before and after the Ag injection. (**G**) Effects of niacin on IgE-dependent PCA. (**H**) The EP3 antagonist attenuated the suppressive effects of niacin on PCA. Tukey’s multiple comparison test was used in all statistical analyses in Fig. 8. Data were pooled from 2 (**A**, **G**, **H**), 3 (**D**), or 4 (**B**) independent experiments. *, *p*<0.05.

### Effects of the niacin treatment on anaphylaxis

To elucidate the roles of the GPR109A stimulation on IgE-mediated anaphylaxis, we evaluated the effects of niacin on PCA. Niacin is general term for two kinds of vitamin B3, nicotinic acid and nicotinamide. Although both compounds similarly plays important roles in nutrition as vitamin B3, nicotinic acid and nicotinamide are distinguishable on the basis of GPR109A stimulation activity, namely, nicotinic acid stimulates GPR109A, but nicotinamide does not. As shown in Fig. 8G, nicotinic acid significantly reduced footpad swelling induced by PCA, whereas nicotinamide did not. The suppression of PCA by the nicotinic acid treatment was inhibited by the additional administration of NSAIDs (data not shown) or the EP3 antagonist (Fig. 8H).

These results demonstrated that the stimulation of GPR109A was involved in the suppression of anaphylaxis by modulating the production of prostanoids, particularly PGE_2_.

## Discussion

MCs play important roles in IgE-dependent allergic responses. The cross-linking of IgE-binding FcεRI on MCs via allergens induces rapid responses, including degranulation and the release of eicosanoids, and late-phase inflammation accompanied by the production of cytokines. Current therapeutic medicines for allergies, such as histamine receptor antagonists, leukotriene receptor antagonists, and humanized anti-human IgE, target steps in the IgE-MC axis, suggesting that the inhibition of MC activation is useful for the prevention of allergic diseases.

In the present study, we identified GPR109A-mediated signaling as a key event for the suppression of MC-mediated allergic responses. A recent study demonstrated that GPR109A was involved in colonic homeostasis based on a *Gpr109a* gene deficiency exacerbating dextran sodium sulfate (DSS)-induced colonic inflammation ^28^. Furthermore, the aforementioned study indicated that the activation of GPR109A by niacin or butyrate suppressed colitis and colon cancer, and colonic macrophages, dendritic cells (DCs), and the epithelium were considered to be the main target cells expressing GPR109A. GPR109A signaling amplified the production of retinoic acid and anti-inflammatory cytokines from macrophages and DCs and subsequently accelerated the development of Tregs. The involvement of butyrate in CD103^+^ DC-mediated Treg differentiation was supported by another study using a food allergy mouse model with high-fiber feeding ^29^. High-fiber feeding enhanced retinaldehyde dehydrogenase (RALDH) activity in CD103^+^ DCs in a GPR109A-dependent manner. As reported in previous studies, DCs and macrophages were identified as GPR109A-expressing myeloid cells. In contrast, the role of GPR109A in MCs was largely unknown. In the present study, we found that *GPR109A* mRNA levels were higher than those of *GPR43* (and *GPR41*) in BMMCs and human MCs (Figs. 4B, 4E). The knockdown of *Gpr109a* completely and partially canceled the suppressive effects of valerate and butyrate, respectively, and exacerbated the degranulation of control MCs. The overexpression of GPR109A reduced the degranulation degree and enhanced the sensitivity of MCs to SCFAs. These results demonstrated the following: i) GPR109A signaling inhibits MC activation, ii) valerate is one of the ligands for GPR109A, and iii) endogenous ligands for GPR109A are present in the MC culture medium. Furthermore, the administration of butyrate, valerate, or niacin prior to passive anaphylaxis significantly ameliorated IgE-dependent anaphylaxis (Figs. 1, and 8). The results of *in vitro* and *in vivo* experiments revealed the potential of the butyrate/valerate/niacin-GPR109A axis as a therapeutic target for MC-mediated allergy. In a previous study, the provision of butyrate in drinking water *ad libitum* for 3 weeks reduced the anaphylaxis score of food allergy in mice, and this was accompanied by a decrease in serum IgE levels and increase in CD103^+^ DCs and the Treg population in MLNs ^29^. Although the effects of butyrate on MC functions were not evaluated ^29^, the oral intake of butyrate or valerate for 1 week suppressed PSA and PCA in the present study (Figs. 1, and 8); therefore, butyrate may have contributed to the decrease observed in the anaphylaxis score with the suppression of MC functions in the previous study. In the present study, Treg levels were not elevated in mice orally administered butyrate or valerate for one week (Fig. 1). Therefore, we concluded that butyrate and valerate ameliorated anaphylaxis by directly modulating MC functions even when Treg development was not affected by SCFAs.

The protective effects of niacin against colitis were also reported in a study using a DSS- or 2,4,6-trinitrobenzene sulfonic acid (TNBS)-induced mouse model ^30^. The administration of niacin up-regulated PGD_2_ levels in the colon and subsequently ameliorated DSS/TNBS-induced colitis in mice via the DP1 receptor ^30^. MCs are a major source of PGD_2_. A previous study using mice with food allergy revealed an anti-allergic role for PGD_2_ derived from MCs ^31^. However, under our experimental conditions, PGE_2_, but not PGD_2_, suppressed MC activation in an autocrine manner (Fig. 7). Another study using ptges^-/-^ mice demonstrated that a PGE_2_ deficiency caused aspirin-exacerbated respiratory diseases (AERD) in which overproduced cysteinyl leukotrienes (cysLTs) and activated MCs were involved in airway inflammatory disorders ^32^. PGE_2_ have been shown to attenuate the cysLT-mediated IL-33-dependent activation of MCs ^32, 33^, whereas MC-derived PGD_2_ is one of the hallmarks of the severity of AERD ^34, 35^. Furthermore, PGE_2_ exhibited immunosuppressive effects in the colonic mucosa ^36^. SCFAs, which enhanced PGE_2_ production by MCs (Fig. 7), may contribute to the prevention and/or treatment of AERD and inflammatory bowel disease.

A recent study reported that PGE_2_ levels inversely correlated with the severity of anaphylaxis in humans and mice ^37^. In that study, an EP2 agonist and EP4 agonist suppressed the IgE-induced activation of BMMCs and IgE-mediated PSA. In contrast, the EP3 antagonist, but not the EP4 antagonist inhibited the protective effects of butyrate and niacin against anaphylaxis under our experimental conditions (Fig. 8). Although we cannot explain this discrepancy, EP3 plays protective roles in MC-mediated allergic responses because *Ptger3^-/-^* mice had more severe airway inflammation accompanied by the hyperactivation of MCs ^38^.

The characteristics of butyrate and valerate differed in the present study. For example, TNF-α production by MCs was decreased by valerate and increased by butyrate (Fig. 2). Valerate primarily suppressed MC activation via GPR109A, whereas butyrate affected multiple pathways in MCs, including those associated with other GPCRs, in addition to GPR109A (Fig. 4). Two other GPCRs, GPR41 and GPR43, which are receptors for SCFAs ^2–4, 6^, may be involved in MC functions. However, we were unable to clarify this aspect because siRNAs for GPR41 and GPR43 were not functional in MCs, in which mRNAs for GPR41 and GPR43 were rarely detected. Furthermore, a previous study demonstrated that the deficiencies in GPR41 and GPR43 in mouse MCs did not affect the butyrate-mediated suppression of degranulation ^9^. An observation that acetate, whose specificity is restricted to GPR43 (EC_50_ for acetate is ∼250-500 μM) and barely to GPR41 (propionate > butyrate >> acetate) but not to GPR109A ^39^, dose not suppress IgE-mediated activation of MCs (^9^ and Fig. 2), may support the irrelevance of GPR41 and GPR43 in the activation of MCs.

*In vivo* experiments using TSA revealed that HDACi, which is one of the activities of butyrate and valerate, inhibited the IgE-induced degranulation and cytokine release of MCs (Fig. 5). The suppressive effects of HDACi on the IgE-dependent activation of MCs were reported in previous studies, in which the down-regulation of tyrosine kinases ^9^ and the suppression of transcription factor recruitment ^40^ were observed. In the present study, the cell surface expression level of FcεRI was decreased by both SCFAs and TSA (Figs. 3, and 5). Although the down-regulation of BTK, SYK, and LAT in butyrate-treated MCs was transcriptional due to a decrease in the acetylation of these promoters ^9^, the mRNA levels of the three subunits of FcεRI slightly increased in SCFA-treated MCs (Fig. 3) and TSA-treated MCs (Fig. 5). Further studies are warranted to clarify the mechanisms by which SCFAs reduce the surface expression of FcεRI on MCs via HDACi activity.

The composition of the daily diet has been shown to affect the microbiota population and SCFA concentrations, which are closely associated with human and animal health ^41, 42^. In contrast to acetate, butyrate, and propionate, which are major SCFAs in the colon, limited information is currently available on valerate, isobutyrate, and isovalerate, the colonic concentrations of which are low. *Clostridia* and Bacteroides have the potential to produce valerate ^41, 43, 44^. Isobutyrate and isovalerate, which suppressed MC activation (Fig. 2), are produced as secondary metabolites from amino acids in *Bacillus*^45^. Both of these branched SCFAs are present in the Japanese traditional food “Natto”, which is made of soybeans through a fermentation process by *Bacillus subtilis natto*. Although branched SCFAs in Natto have been recognized merely as the source of a unique flavor, they may exert anti-allergic effects and/or contribute to colonic homeostasis. Changes in dietary habits in Japan in the past few decades, from a typical Japanese diet including Natto and high-fiber dishes to Western diets may be associated with the incidence of allergic diseases. We intend to investigate the relationship between the mucosal environment and immunoregulation by focusing on valerate, butyrate, and branched SCFAs, which may lead to the proposal of the health benefits of certain foods in the near future.

## Methods

### Mice and cells

BMMCs were generated from the BM cells of C57BL/6 mice (Japan SLC, Hamamatsu, Japan) by cultivation in the presence of 5 ng/mL of mouse IL-3 (BioLegend) as previously described ^18, 46, 47^. All animal experiments were performed in accordance with the approved guidelines of the Institutional Review Board of Tokyo University of Science, Tokyo, Japan. The Animal Care and Use Committees of Tokyo University of Science approved this study (K22005, K21004, K20005, K19006, K18006, K17009, K17012, K16007, K16010).

### Reagents

SCFAs (acetate, butyrate, sodium butyrate, isobutyrate, propionate, valerate, ethylvalerate, and isovalerate), a monocarboxylate transporter inhibitor (α-cyano-4-hydroxycinnamic acid, CHCA) (Cat. 476870), and acetylsalicylic acid (ASA) (Cat. A5376) were purchased from Sigma-Aldrich (St. Louis, MO). Pertussis toxin (PTX), which is used as a Gi/o-type G protein inactivator, was purchased from Calbiochem (Cat. 516561, San Diego, CA), and indomethacin was obtained from Fujifilm Wako Pure Chemical (Cat. 095-02472, Japan). PGD_2_ (Cat. 12010), PGE_2_ (Cat. 14010), and an inhibitor of COX-1, SC-560 (CAY-70340) were supplied by Cayman Chemical (Ann Arbor, MI). Nicotinic acid/niacin (#72309, Sigma-Aldrich) and nicotinamide/niacinamide (#N0078, Tokyo Chemical Industry Co., Ltd., Tokyo, Japan) were obtained. ONO-AE5-599, an antagonist of PGE_2_ receptor 3 (EP3), and ONO-AE3-208, an antagonist of EP4 were kindly provided by Ono Pharmaceutical Co. Ltd. (Osaka, Japan)

### Quantification of mRNA

The purification of total RNA and reverse transcription to synthesize cDNA were performed using a ReliaPrep RNA Cell Miniprep System (Promega) and ReverTra Ace qPCR RT Master Mix (TOYOBO, Osaka, Japan), respectively. The mRNA levels were measured by quantitative PCR using a Step-One Real-Time PCR system (Applied Biosystems) with THUNDERBIRD probe qPCR Mix (TOYOBO) or THUNDERBIRD SYBR qPCR Mix (TOYOBO). The TaqMan primers *Gata1* (#Mm00484678_m1), *Gata2* (#Mm00492300_m1), *Spi1* (#Mm01270606_m1), *Fcer1a* (#Mm00438867_m1), *Ms4a2* (#Mm00442780_m1), *Fcer1g* (#Mm00438869_m1), and *Gapdh* (#4352339E) were purchased from Applied Biosystems. The following oligonucleotides were synthesized for PCR primers: *Gpr41* forward, 5’-GTGACCATGGGGACAAGCTTC-3’, reverse, 5’-CCCTGGCTGTAGGTTGCATT-3’, *Gpr43* forward, 5’-GGCTTCTACAGCAGCATCTA-3’, reverse, 5’-AAGCACACCAGGAAATTAAG-3’, *Gpr109a* forward, 5’-ATGGCGAGGCATATCTGTGTAGCA-3’, reverse, 5’-TCCTGCCTGAGCAGAACAAGATGA-3’, *Slc5a8* forward, 5’-CATTCGTCTCTGTGGCACAATC-3’, reverse, 5’-GGGCATAAATCACAATTCCAGTGT-3’, *Slc5a12* forward, 5’-ACAACAACAGTAGCCCCACAGA-3’, reverse, 5’-GGTAGGAGAGTGAGTACCATGTGTCA-3’, Slc16a1 forward, 5’-ACAACAACAGTAGCCCCACAGA-3’, reverse, 5’-GGTAGGAGAGTGAGTACCATGTGTCA-3’.

### Degranulation assay

Two hundred nanograms of anti-TNP mouse IgE (clone IgE-3, BD Bioscience, San Jose, CA) was added to 1 mL of culture medium containing BMMCs (5 × 10^5^). After a 2-h incubation, cells were washed with Tyrode’s buffer and resuspended in Tyrode’s buffer containing TNP-BSA (final concentration of 3 ng/mL) (LSL, Tokyo, Japan). Thirty minutes after the TNP-BSA stimulation, the culture supernatant was harvested, and β-hexosaminidase activity in the supernatant was assessed as previously reported ^17^.

### Flow cytometry

Cell surface FcεRI and c-kit stained by PE-labeled anti-mouse FcεRIα Ab (MAR-1, eBioscience) and FITC-labeled anti-mouse CD117 Ab (2B2, BioLegend) were detected by a MACS Quant (Miltenyi Biotech). To detect Foxp3^+^ cells in CD4^+^ T cells, cells treated with a Foxp3/Transcription factor staining buffer kit (Cat. TNB-0607, TOMBO Bioscience) were stained with FITC-labeled anti-mouse CD4 Ab (GK1.5, BioLegend) and allophycocyanin-labeled anti-mouse Foxp3 Ab (3G3, TOMBO Bioscience).

### Knockdown by small interfering RNA (siRNA)

siRNA for mouse Gpr109a (MSS234551), and control siRNA (Stealth RNAi Negative Universal Control, #12935) were purchased from Invitrogen (Carlsbad, CA). Ten microliters of 20 μM siRNA was introduced into BMMCs (5 × 10^6^) by a Neon Transfection System (Invitrogen) set at Program #5 using a Neon 100 μL Kit.

### Overexpression of mouse GPR109A on MCs by a retroviral vector

Mouse GPR109A cDNA was amplified by PCR using mouse BM macrophage cDNA as a template, and the following oligonucleotides as primers: 5’-ggc*ggatcc*<u>atg</u>agcaagtcagaccattttctag-3’ (the inserted *Bam*HI sequence is shown in italics, and the initiation codon is underlined) and 5’-ggg*ctcgag*<u>tta</u>acgagatgtggaagccag-3’ (the inserted *Xho*I site and termination codon are shown in italics and underlined, respectively). The *Bam*HI/*Xho*I-digested PCR fragment encoding GP109A cDNA was inserted into the *Bam*HI/*Xho*I region in the multicloning site of pMXs-puro ^48^ to obtain pMXs-puro-mGPR109A. The preparation of retroviral vectors using pMX plasmids and packaging cells, the transfection of BMMCs by retroviral vectors, and the selection of transfectants were performed as previously described ^49, 50^.

### ELISA

The concentrations of IL-13 and TNF-α in the culture media were measured by using mouse IL-13 DuoSet ELISA (R&D Systems, Minneapolis, MN) and ELISA MAX Deluxe Set mouse TNF-α (BioLegend), respectively. Prostaglandin E_2_ ELISA Kit-Monoclonal (Cayman Chemical, # 514010) was used to determine the concentration of prostaglandin E_2_ (PGE_2_).

### Passive systemic anaphylaxis (PSA) and the oral administration of SCFAs

C57BL/6 mice were orally administered 435 mg/kg/day butyrate or 510 mg/kg/day valerate for 4 to 6 days using saline and ethanol-PEG as a vehicle. On the last day of the administration, mice received an intravenous (i.v.) injection of 3 μg of TNP-specific mouse IgE. Twenty-four hours after the IgE injection, mice were i.v. transfused with 20 μg of TNP-BSA. The body temperature of each mouse was measured every 10 min for 1 h.

### Passive cutaneous anaphylaxis (PCA)

After 4 days of the administration of butyrate, 0.02 μg of TNP-specific mouse IgE or saline as a control was injected into the footpad to establish PCA. Twenty-four hours after the IgE injection, mice were i.v. transfused with 20 μg of TNP-BSA. Footpad thickness was measured 0.5 h after the TNP-BSA injection.

### Administration of ASA, indomethacin, an EP3 antagonist, an EP4 antagonist, niacin/nicotinic acid, and niacinamide

Mice were orally administered ASA (25 mg/kg), indomethacin (1 mg/kg), ONO-AE5-599 (10 mg/kg), ONO-AE3-208 (10 mg/kg), or vehicle (0.5 % methyl cellulose, 0.2 mL) twice per a day using a disposable feeding needle with a 200-μL scale (#7202, Fuchigami Co., Ltd., Kyoto, Japan). Niacin/nicotinic acid (100 mg/kg), niacinamide (100 mg/kg), or vehicle (saline, 0.2 mL) was administered via an intraperitoneal (i.p.) injection once per a day.

## Statistical analysis

A two-tailed Student’s *t*-test was used to compare two samples, and a one-way ANOVA-followed by Tukey’s multiple comparison test, Dunnett’s multiple comparison test, and Sidak’s multiple comparison test were used to compare more than three samples. *p* values < 0.05 were considered to be significant.

## Acknowledgments

We are grateful to the members of the Laboratory of Molecular Biology and Immunology, Department of Biological Science and Technology, Tokyo University of Science, for their constructive discussions and technical support. This work was supported by a Grant-in-Aid for Scientific Research (B) 20H02939 (CN); Grants-in-Aid for Scientific Research (C) 21K05297 (MH), 19K05884 (TY), and 19K08920 (KK); a Research Fellowship for Young Scientists DC2 and a Grant-in-Aid for JSPS Fellows 21J12113 (KN); a Scholarship for Doctoral Student in Immunology (from JSI to NI); a Tokyo University of Science Grant for President’s Research Promotion (CN); the Tojuro Iijima Foundation for Food Science and Technology (CN); a Research Grant from the Mishima Kaiun Memorial Foundation (CN); and a Research Grant from the Takeda Science Foundation (CN).

## Authorship contribution

K.N. performed the experiments and wrote the manuscript; D.A. performed the experiments and analyzed data; T.A., K.I., R.M., I.F., M.A., N.I., H.K., Y.I., M.I., and T.Y. performed the experiments; M.H. analyzed data and wrote the manuscript; K.K. designed the research and performed the experiments; C.N. designed the research and wrote the manuscript.

## Additional information

Supplemental information includes two figures.

## Abbreviations used

AERD: aspirin-exacerbated respiratory diseases
Ag: antigen
ASA: acetylsalicylic acid
BM: bone marrow-derived
cysLT: cysteinyl leukotriene
DC: dendritic cell
GPCR: G protein-coupled receptor
HDAC: histone deacetylase
i.p.: intraperitoneal
i.v.: intravenous
MC: mast cell
NSAID: non-steroidal anti-inflammatory drug
PCA: passive cutaneous anaphylaxis
PGs: prostaglandins
p.o.: per os
PSA: passive systemic anaphylaxis
PTX: pertussis toxin
RALDH: retinaldehyde dehydrogenase
s.c.: subcutaneous
SCFA: short chain fatty acid
siRNA: small interfering RNA
TSA: trichostatin A.

