## Supplementary material for "Butyrate, valerate, and niacin ameliorate anaphylaxis by suppressing IgE-dependent mast cell activation: Roles of GPR109A, PGE_2_, and epigenetic regulation": Nagata-Ando-bioRxiv-supplemental.docx

**Butyrate, valerate, and niacin ameliorate anaphylaxis by suppressing IgE-dependent mast cell activation: Roles of GPR109A, PGE_2_, and epigenetic regulation in anti-allergic effects of short chain fatty acids**

Running title: Effects of SCFAs in IgE-dependent MC activation

**Kazuki Nagata^1^, Daisuke Ando^1^, Tsubasa Ashikari^1^, Kandai Ito^1^, Ryosuke Miura^1^, Izumi Fujigaki^1^, Miki Ando^1^, Naoto Ito^1^, Hibiki Kawazoe^1^, Yuki Iizuka^1^, Mariko Inoue^1^, Takuya Yashiro^1^, Masakazu Hachisu^1^, Kazumi Kasakura^1^, and Chiharu Nishiyama^1*^**

^1^ Department of Biological Science and Technology,

Faculty of Advanced Engineering, Tokyo University of Science,

6-3-1 Niijuku, Katsushika-ku, Tokyo 125-8585, Japan

K.N. and D.A. contributed equally to this work.

**Supplementary Figures and Figure Legends**

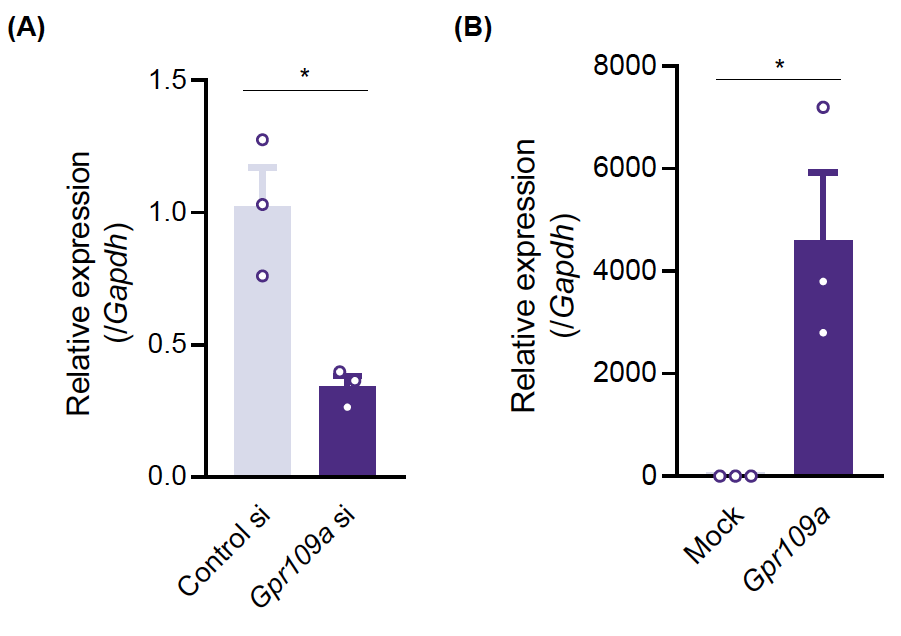

**Figure S1. mRNA levels of *Gpr109a* in MCs with the knockdown or overexpression of GPR109A.**

(**A**) BMMCs introduced *Gpr109a* siRNA or control siRNA were harvested 48 h after transfection to measure mRNA levels, and were used for degranulation assay (**Figure 4F**).

(**B**) BMMCs transfected with retrovirus carrying *Gpr109a* cDNA or control virus were harvested puromycin selection for 10 days, and were used for degranulation assay (**Figure 4G**).

Data represent the mean ± SEM of 3 independent experiments. A two-tailed Student’s *t*-test was used for statistical analysis. *, *p*<0.05.
